# Commercial fishery disturbance of the global open-ocean carbon sink

**DOI:** 10.1101/2020.09.21.307462

**Authors:** E. L. Cavan, S. L. Hill

## Abstract

Primary production in the global oceans fuels multiple ecosystem services including fisheries, and the open-ocean biological carbon sink, which support food security and livelihoods^1^, and the regulation of atmospheric CO_2_ levels^2^ respectively. The spatial distributions of these two services are driven by primary production and it is likely that ecosystem disturbance from fishing impacts both the carbon sink and atmospheric CO_2_. Yet the extent of these impacts from past, present and future fishing is unknown. Here we show that 23% of global export and 40% of fishing effort are concentrated in zones of intensive overlap representing 7% of the global ocean area. This overlap is particularly evident in the Northeast Atlantic and Northwest Pacific. Small pelagic fish dominate catches in these regions and globally, and their exploitation will reduce faecal pellet carbon sinks and may cause tropic cascades affecting plankton communities important in sinking carbon. There is an urgent need to address how fisheries affect carbon cycling, and for policy objectives to include protecting the carbon sink, particularly in areas where fishing intensity and carbon export and storage are high.

---

The open-ocean carbon sink and store via the biological pump^2,3^, hereafter ‘carbon sink’, is an important regulator of atmospheric CO_2_ levels, which would otherwise be 50 % higher^4^. Estimates of organic carbon exported out of the top 100 m of the global ocean range from 4 – 12 Gt C yr^-1^ ^5,6^. Exported carbon sinks down to the deep ocean (> 1000 m) where ~ 1 % is locked away on timescales from decades to millennia, with the rest being recycled and eventually converted back to CO_2_ by microbes and zooplankton^3^. This 1 % of carbon export equates to deep ocean carbon sequestration of up to 0.12 Gt C yr^-1^, which is on a par with coastal blue carbon sequestration (0.11 Gt C yr^-1^ from mangroves, salt marshes and seagrass^7^), or 1.1 % of anthropogenic carbon release (10 Gt C yr^-1^)^8^. The open-ocean carbon sink is predominantly driven by phyto- and zooplankton at the base of ocean food-webs^3^. The faecal pellets of current and potential fishery species, including anchovy^9^, krill^10^ and mesopelagic fish^11^, are particularly important in sinking. Any marine ecosystem change resulting in changes in abundance or community composition of species responsible for sinking and storing carbon could result in a positive feedback increasing atmospheric CO_2_ levels^12^.

Marine fishing currently removes ~ 0.10 Gt yr^-1^ of biomass^13^ and has profoundly altered ecosystems throughout the global ocean. These impacts can propagate through foodwebs in trophic cascades which produce sequential changes in the abundance of successive trophic levels^14^. Fishing also affects the physical habitat, such as through the removal of oyster beds^15^. These ecological alterations can affect the lower trophic levels responsible for the majority of carbon fixation and export, and those that contribute to deeper faecal carbon sinks. The reliance of both fish biomass and the carbon sink on phytoplankton^1,2^ creates the potential for significant spatial overlap between the two and for the fishing to disturb the carbon sink. Although there is some acknowledgement of potential interactions between the two^16^, the impact of past and current fishing on the carbon sink and atmospheric CO_2_ has not been investigated, nor is fishery disturbance factored in to forecasts of future changes to the global carbon cycle^17^.

The main reason for the lack of attention to this topic is likely a discipline divide between biogeochemistry and marine ecology. This divide is reflected in models; the biogeochemical modules of the Earth System Models (ESMs) which inform Intergovernmental Panel on Climate Change (IPCC) assessment reports do not include trophic levels above zooplankton^18^. While ecological modellers are working to better link ESMs and models of fished species^19^, the primary motivation is to investigate the bottom-up impacts of climate change on these species^20^, rather than top-down controls on the global carbon sink.

The current study uses global scale satellite data to assess the spatial overlap between commercial fishing effort^21^ and the carbon sink (specifically particulate carbon export at 100 m depth)^6^, thereby mapping the risk of impact. We analyse these data at two scales, namely a 1° x 1° grid and the nineteen major fishing areas (hereafter ‘fishing area’) used by the UN Food and Agricultural Organisation (FAO) for recording catch statistics. We also identify the routes by which different fishing practices might impact the carbon sink.

## Regions of high carbon sink and fishing

Both carbon export and fishing intensity are highest around coastlines (Fig. 1), which is reflected in the map showing areas of combined high carbon export and high fishing intensity (Fig. 2). Both ecosystem services are concentrated in coastal regions where primary production is highest^22^. The spatial overlap (orange pixels in Fig. 2) represents 7% of the global oceans by area, but 23% of carbon export and 40 % of fishing effort globally. The two highest ranking areas, for both carbon export and fishing intensity, are the Northeast Atlantic (fishing area 27, Fig. 2) and the Northwest Pacific (fishing area 61). These areas are respectively responsible for 14% and 9% (0.46 and 0.32 Gt C yr^-1^) of global carbon export and 15% and 14% (33.26 x 10^6^ and 29.99 x 10^6^ hours yr^-1^) of global fishing effort (Fig. 3, Supplementary Table 1).

**Fig. 1.**
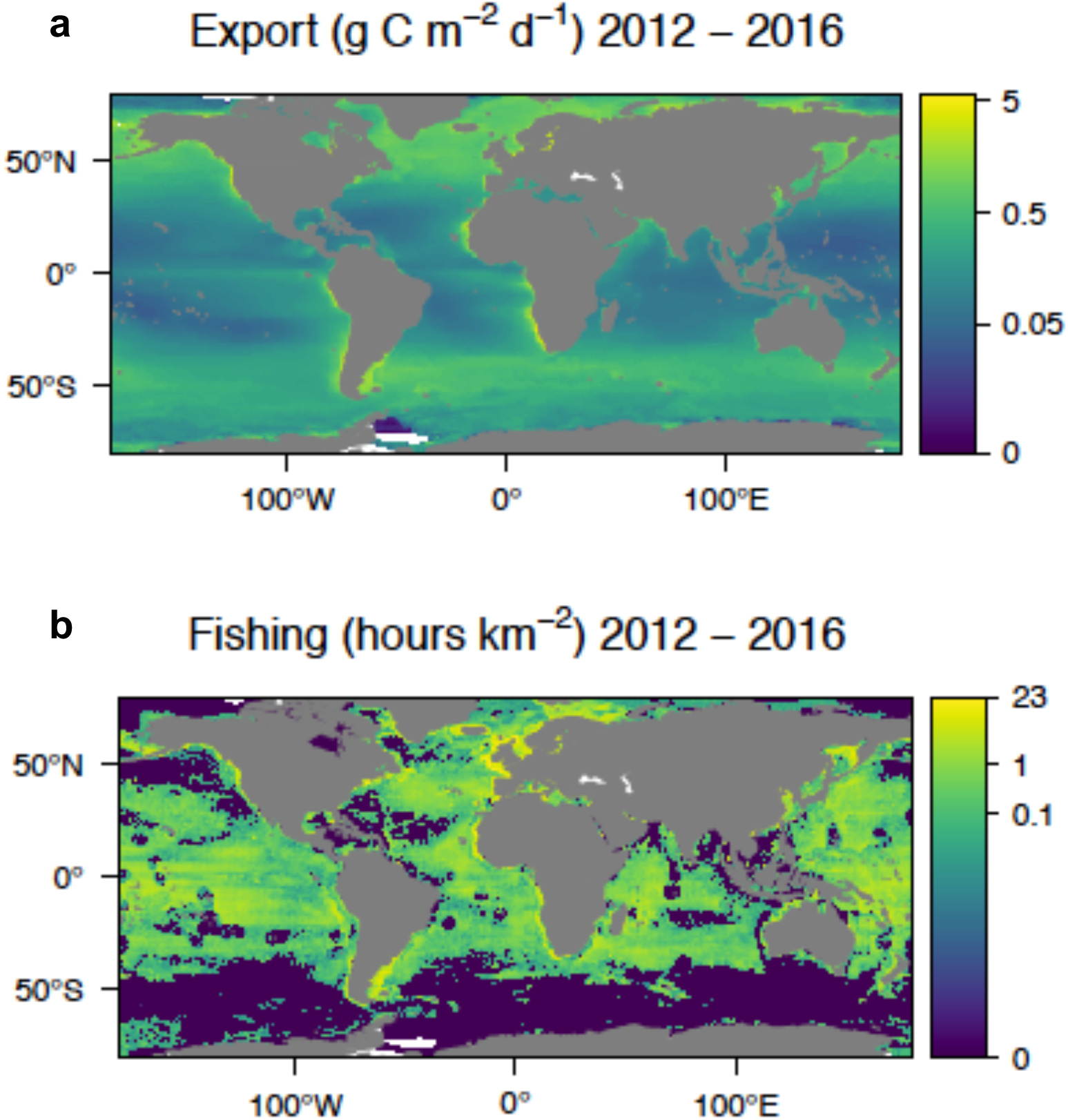
Global annual carbon sink (export) and fishing intensity. **a)** Average annual particulate organic carbon export (g C m^-2^ d^-1^) from 100 m depth estimated using satellite primary production and sea surface temperature according to the algorithm in Henson et al^6^. b) Average annual commercial (vessels 6 – 146 m in length) fishing intensity (hours fished km^-2^), data downloaded from Global Fishing Watch^21^. Both datasets are averaged over a 5-year period from 2012 – 2016, note the log z-scale.

**Fig. 2.**
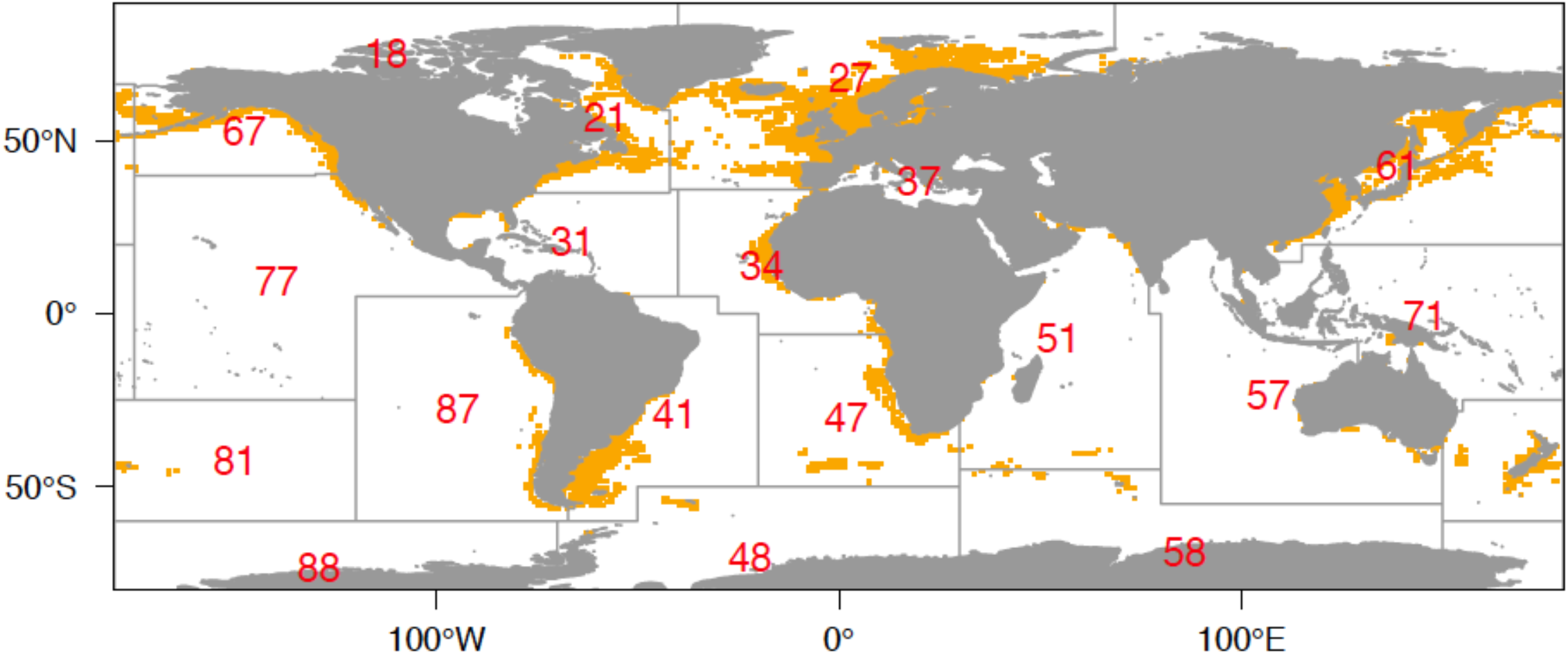
Regions of high fishing and carbon export intensity. 1° by 1° grid cells where carbon export (Fig. 1a) and fishing hours (Fig. 1b) values are in the upper quartile of both data sets, emphasising the importance of coastal regions at higher latitudes, particularly the Northeast Atlantic (fishing area 27) and Northwest Pacific (fishing area 61) (see Fig. 3 and Supplementary Table 1). Grey grid lines and red numbers indicate the FAO major fishing areas.

**Fig. 3.**
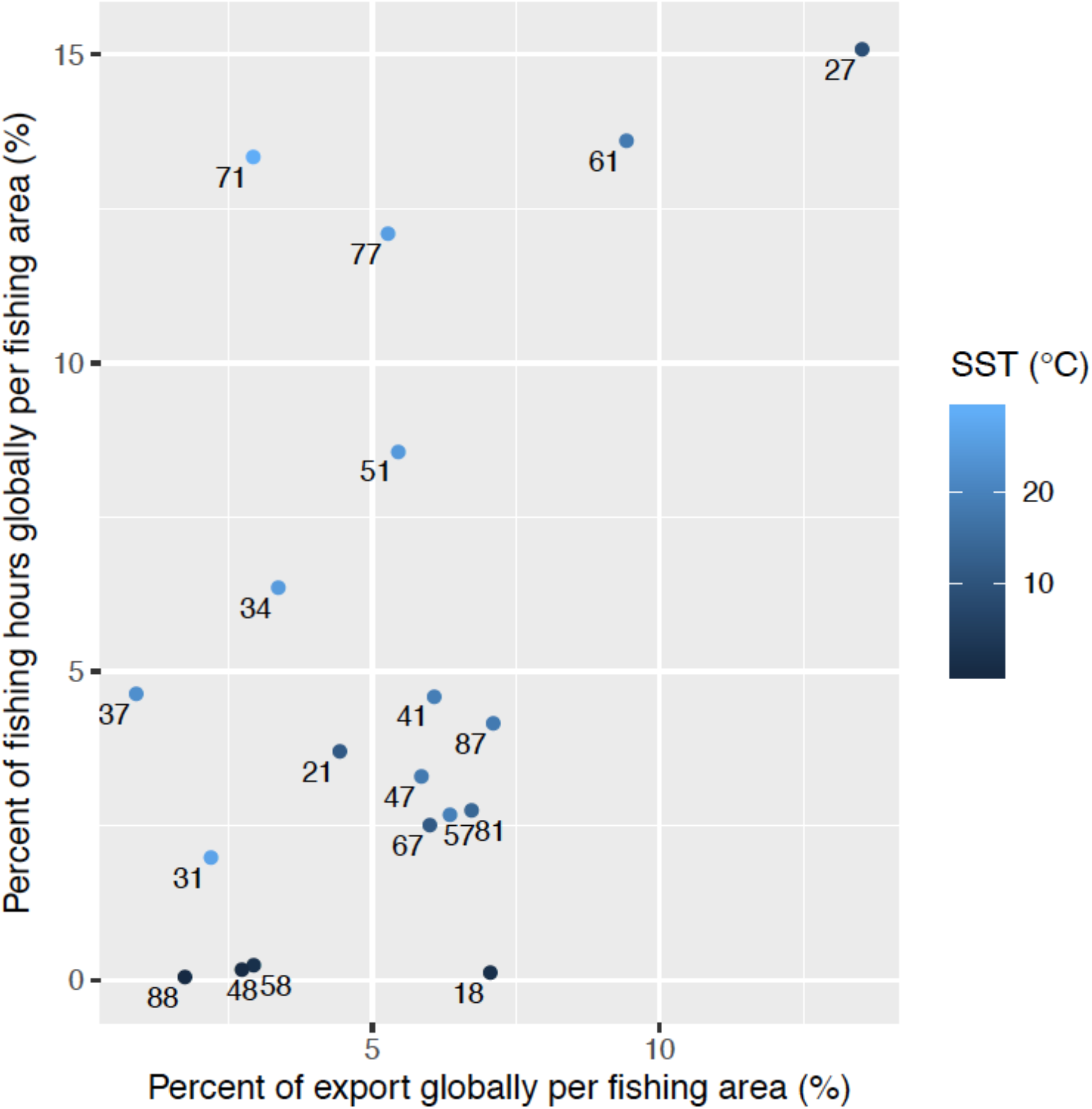
Relationship between carbon export and fishing intensity across fishing areas. Percent of global particulate organic carbon export and fishing intensity in each fishing area averaged over 2012 - 2016. Colour of points present the mean sea surface temperature (SST) of each fishing area (Supplementary Table 1) and the labels refer to fishing area number. Fishing areas 27 (Northeast Atlantic) and 61 (Northwest Pacific) have highest carbon export and fishing intensity. Fishing area 18 is the Arctic where fishing is minimal, but the export is relatively high due to high primary production and low temperatures.

Fishing intensity increases with total carbon export at the fishing area scale (Fig. 3). The Arctic fishing area (18) does not follow this pattern as it has high carbon export but relatively little fishing effort due to seasonal ice cover, although melting sea ice may change this in the future^23^. Subtropical fishing areas (Central West Pacific, 71, Central East Pacific, 77, and West Indian, 51) have high total fishing intensity (13 %, 12 % and 9 % of global total respectively), but fairly low carbon export (≤ 5 %) (Fig. 3). Fishing areas which contain coastal upwelling regions (e.g. Southeast Pacific, 87, and Southeast Atlantic, 47) make relatively low contributions to global fishing and export (Fig. 3) because they are dominated by low productivity oligotrophic gyres (Fig. 1a). The high localised primary production, and thus carbon export and fishing, in coastal upwellings is nonetheless highlighted by our upper quartile analysis of both data sets (Fig. 2).

## Impacts of fishing on the carbon sink

From our analysis of FAO catch data, we identified small and medium (< 60 cm length, hereafter small) pelagic fish as the dominant fished group globally, with trawls the dominant gear type. In the Northeast Atlantic where fishing intensity and carbon export are highest, Atlantic mackerel and Atlantic herring dominate the catch, and in the Northwest Pacific Japanese anchovy is the main fished small pelagic. Fishing small pelagics can have both direct and indirect impacts on the carbon sink. These fish contribute to the carbon sink through releasing carbon-rich and fast sinking faecal pellets that can sink at > 700 m d^-1^ ^9^ (Fig. 4). For example Peruvian anchoveta may be responsible for around 7 % of local carbon export^24^. Reducing the biomass of these species will reduce the carbon faecal pellet sink, which is one of the most important routes to sink organic carbon^25^. Whether the removal of small pelagics indirectly impacts the lower trophic levels through trophic cascades remains uncertain. Cod fishing in the Baltic Sea increased small pelagic (sprat) biomass, which led to a reduction in its zooplankton prey as part of a more extensive trophic cascade^26^ (Fig. 4). However, specific evidence of the existence or extent of indirect impacts caused by trophic cascades is lacking for major fished species, including for Atlantic herring, mackerel and Japanese anchovy

**Fig. 4.**
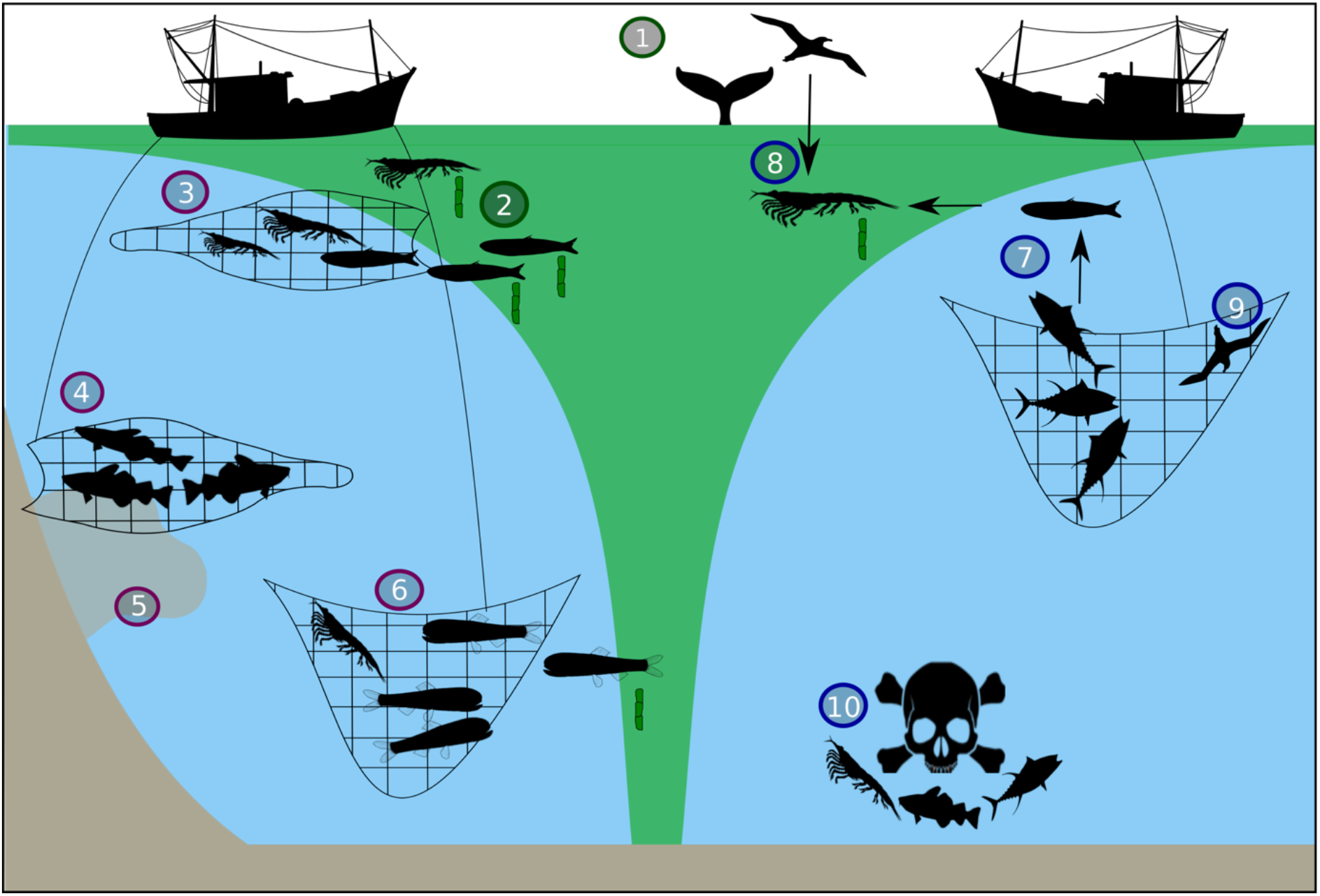
Direct and indirect impacts of fishing to the carbon sink. Phytoplankton (green shading in the surface) stimulate fish biomass production and the export of carbon out of the upper ocean, of which ~ 1 % sinks to the deep ocean. The carbon sink is enhanced by (1) fertilising species and (2) those egesting fast-sinking carbon-rich faecal pellets. Direct impacts of fishing include (3) harvesting low-mid trophic level pellet-producing species, (4) removing species living near the seabed where the sink of carbon will be short, and (5) harvesting groundfish disturbing the sediment resuspending carbon which could be remineralised in the water to CO_2_, and finally (6) removing resident or migratory mesopelagic species that contribute to the carbon sink. Indirect impacts include (7) causing trophic cascades when removing high trophic level species impacting low trophic level communities that sink carbon, (8) removing prey items for fertilizing species (e.g. mackerel or krill that feed seabirds), (9) killing predators (e.g. seabirds) that may otherwise fertilise the oceans but also help to maintain a balanced food web, and finally (10) the release discards which could cause localized dead zones.

Groundfish such as Atlantic cod and Alaska pollock (caught in fishing areas 27 and 61 respectively) are the next most important catch category after small pelagics (Supplementary Table 1), but their contribution to the carbon sink is also currently unknown. Groundfish fisheries could have the greatest impacts on the carbon sink through trophic cascades as described above in the Baltic Sea^26^ and physical disturbance of the seabed^27,28^ (Fig. 4). The demersal trawls used in these fisheries create plumes of resuspended material that can remove seabed carbon at a rate that counteracts sinking carbon^28^. As groundfish reside near the seabed, the pellets they egest would be subjected to less water column degradation prior to sedimentation of the carbon. Similarly, mesopelagic fish that live permanently or migrate daily into this depth realm can increase the sink of carbon to the deep sea and seabed^11^; any carbon they release below the permanent thermocline (winter mixed layer depth) will not be subject to water column mixing and remain sequestered for decades or centuries^10^. Thus targeted or incidental harvesting of mesopelagic species is likely to increase the rate at which CO_2_ returns to the atmosphere (Fig. 4). Other mechanisms by which fishing for any species could impact the carbon sink include the harvesting or by-catch of fertilising species such as krill^29^, whales^30^ or seabirds^31^, and the release of discards causing localized dead zones (see Supplementary Information) or re-routing carbon through different trophic cycles e.g. through scavenging seabirds^32^ (Fig. 4).

## Climate change, fishing and the carbon sink

Global carbon export is projected to decline by the end of the century^33^, as a result of changes to plankton abundance and composition, and reduced primary production^34^. There are no forecasts of how climate change impacts to higher trophic levels will affect the future carbon sink. Fishing may further exacerbate the projected climate-driven declines in carbon export, and thus the store of carbon in the deep ocean, by changing the community composition of low trophic levels important in carbon export. For instance 30 years of warming in the Baltic Sea changed the dominant copepod species from the larger *Pseudocalanus acuspes* to the smaller *Acartia spp*, with overfishing of cod amplifying this regime shift^35^. Climate change will also likely alter the spatial overlap of fishing and carbon export (Fig. 2). Climate-induced spatial shifts have already been observed in fish, including poleward shifts as sea temperatures rise^36^. As for the carbon sink, projections suggest an expansion of oligotrophic regions where carbon export is currently low (Fig. 1a)^37^, and increases in carbon export toward the poles. Poleward shifts in both fishing intensity and the carbon sink would result in smaller, more concentrated areas of overlap than today (Fig. 2), with an increased risk of impact.

## Conclusions

There is clear spatial overlap between the carbon sink and commercial fishing. Biomass and ecosystem changes caused by fishing could negatively impact carbon sinking and storage throughout the water column and seabed, and therefore atmospheric CO_2_ levels. There is an urgent need to clarify through observations and modelling whether and how fisheries reduce the carbon sink, and for policy objectives to include protecting this ecosystem service. These needs are particularly important in the regions where the risk of fishing impacting the carbon sink is high (Northeast Atlantic and Northwest Pacific). Research is also required into the potentially synergistic impacts of fishing on the carbon sink, and climate change on both fishing and the carbon sink. The rebuilding of impacted ecosystems and stocks would help to reverse impacts on the carbon sink. This rebuilding is already an established fisheries management and sustainable development objective^27,38^ but progress towards this goal is extremely limited and up to 63% of monitored stocks remain in need of rebuilding^27,39^. Recognising that the carbon sink is an additional ecosystem service that requires protection strengthens the case for a holistic approach to managing the oceans^27,40^ and might help to achieve a wider suite of environmental goals. We hope improved understanding of how commercial fisheries disturb the carbon sink will be a step toward realising a sustainable balance of the twin needs for productive fisheries to maintain global food security and strong carbon sinks which play a critical role in climate regulation.

## Methods

Our indicator of carbon sink intensity (export) is the critical first step in the carbon sink while our indicator of fishing (effort) is correlated with the main potential route of impact, i.e. biomass removal (see Supplementary Information). Sea surface temperature was downloaded from the NASA ocean colour database (https://oceancolor.gsfc.nasa.gov) and primary productivity data from the Ocean Productivity site^41^ for the same time frame as availability of fishing data (2012 – 2016), to calculate particulate organic carbon export (g C m^-2^ d^-1^) sink of carbon out of the top 100 m of the ocean) using the Henson et al.^6^ algorithm. Carbon sinks through the entire ocean depths, but only is stored and sequestered on long timescales if it reaches the deep sea (> 1000 m). However, there is not yet a consensus on how to parameterise the transfer efficiency of carbon to the deep due to the many processes which control it, whereas there is a consensus that carbon export out of the upper 100 m is negatively related to temperature^6,42^. Hence, we use carbon export here as our metric from the global carbon sink. We use data on global fishing intensity (hours fished km^-2^) taken from all vessels with an automatic identification system (AIS) and published online by the Global Fishing Watch^21^. Only data for the years 2012 – 2016 inclusive have been released so we present the mean annual fishing intensity over this 5-year period. We merged global fishing intensity and export data onto a 1 x 1 degree resolution grid and identified the areas where both fishing and export were in the top quartile of their respective datasets globally (orange pixels in Fig. 2).

We assessed the total export, fishing intensity and dominant fishing method (gear type) for each of the FAO major fishing areas. We obtained gear type data primarily from Tanocet et al.^43^, which provides total Global Fisheries Landings database^44^ effort by gear type for 2010 to 2014. We obtained catch data for the equivalent period from the FAO Global Capture Production database^13^ (Supplementary Table 2). This period overlaps our export and fishing intensity data (Fig. 1a and b) for three years, 2012 – 2014, and fishery catch and effort data are well correlated (Supplementary Fig. 1). We identified those taxa which dominate the catch in each fishing area (i.e. the top ranking taxa in terms of catch weight, which constitute 50% or the closest value above 50% of the overall catch) (Supplementary Table 1). We assigned each taxon to one of the following categories: small pelagic fish (SP); groundfish (G); large pelagic fish (LP), deep water fish (DF); unspecified fish (UF), pelagic crustaceans (PC); benthic crustaceans (BC), unspecified crustaceans (UC); squid (S); Unspecified molluscs (UM); and finally bivalves (B). See Supplementary Table 2 for more detail on this classification.

Gear type data were not available for fishing areas in the Southern Ocean (fishing areas 88, 48, 58), nor the Northeast Atlantic (fishing area 27). For the Southern Ocean we were able to characterise our catch data by gear type, providing data that is comparable to the majority of other fishing areas. The dominant Southern Ocean fisheries use either longlines to target toothfish (fishing area 58 & 88) or trawls to target Antarctic krill and mackerel icefish (fishing area 48, Supplementary Table 1). In the case of the Northeast Atlantic, gear type data is presented in terms of percentage of fishing hours rather than percentage of catch^43^. Our Supplementary Table 1 presents these data, which suggest that trawls are the main fishing gear in the Northeast Atlantic, comprising more than 70% of fishing hours. It is therefore plausible that trawls are also the main fishing gear by catch, although the two metrics are not strictly comparable.

## Acknowledgements

E.L.C was supported by an Imperial College Research Fellowship funded by Imperial College London. S.H was supported by Natural Environment Research Council core funding to the British Antarctic Survey Ecosystems programme.

## Author Contributions

E.L.C conceived the study and analysed the carbon export and fishing intensity data. E.L.C made the figures. S.H. analysed the catch and gear type data. Both authors contributed equally to the development of ideas and the writing and editing of this manuscript.

## Additional Information

Supplementary Information is available for this paper. Correspondence and requests for materials should be addressed to Emma Cavan at or Simeon Hill at.

## References

1. Pauly, D. & Christensen, V. Primary production required to sustain global fisheries. Nature 374, 255 (1995).

2. Volk, T. & Hoffert, M. Ocean Carbon Pumps: Analysis of Relative Strengths and Efficiencies in Ocean-Driven Atmospheric CO2 Changes. in The Carbon Cycle and Atmospheric CO2: Natural Variations Archean to Present 99–110 (American Geophysical Union, 1985).

3. Turner, J. T. Zooplankton fecal pellets, marine snow, phytodetritus and the ocean’s biological pump. Prog. Oceanogr. 130, 205–248 (2015).

4. Parekh, P., Dutkiewicz, S., Follows, M. J. & Ito, T. Atmospheric carbon dioxide in a less dusty world. Geophys. Res. Lett. 33, (2006).

5. Laws, A., Falkowski, G., Smith, O., Hugh, J. & Mccarthy, J. Temperature effects on export production in the open ocean. Global Biogeochem. Cycles 14, 1231–1246 (2000).

6. Henson, S. A. et al. A reduced estimate of the strength of the ocean’s biological carbon pump. Geophys. Res. Lett. 38, L04606 (2011).

7. Duarte, C. M., Middelburg, J. J. & Caraco, N. Major role of marine vegetation on the oceanic carbon cycle. Biogeosciences 2, 1–8 (2005).

8. Zeebe, R. E., Ridgwell, A. & Zachos, J. C. Anthropogenic carbon release rate unprecedented during the past 66 million years. Nat. Geosci. 9, 325–329 (2016).

9. Saba, G. K. & Steinberg, D. K. Abundance, Composition, and Sinking Rates of Fish Fecal Pellets in the Santa Barbara Channel. Sci. Rep. 2, 716 (2012).

10. Cavan, E. L. et al. The importance of Antarctic krill in biogeochemical cycles. Nat. Commun. 10, 1–13 (2019).

11. Davison, P. C., Checkley, D. M., Koslow, J. A. & Barlow, J. Carbon export mediated by mesopelagic fishes in the northeast Pacific Ocean. Prog. Oceanogr. 116, 14–30 (2013).

12. DeVries, T. et al. Decadal trends in the ocean carbon sink. Proc. Natl. Acad. Sci. 116, 11646 LP – 11651 (2019).

13. FAO. FAO yearbook. Fishery and Aquaculture Statistics 2017. (2019).

14. Carpenter, S. R., Kitchell, J. F. & Hodgson, J. R. Cascading Trophic Interactions and Lake Productivity. Bioscience 35, 634–639 (1985).

15. Grabowski, J. H. & Peterson, C. H. 15 - Restoring Oyster Reefs to Recover Ecosystem Services. in Ecosystem Engineers (eds. Cuddington, K., Byers, J. E., Wilson, W. G. & Hastings, A. B. T.-T. E. S.) 4, 281–298 (Academic Press, 2007).

16. Trebilco, R., Melbourne-thomas, J. & John, A. The policy relevance of Southern Ocean food web structure: Implications of food web change for fisheries, conservation and carbon sequestration. Mar. Policy 115, 103832 (2020).

17. Laufkötter, C. et al. Projected decreases in future marine export production: the role of the carbon flux through the upper ocean ecosystem. Biogeosciences Discuss. 12, 3731–3824 (2015).

18. Yool, A., Popova, E. E. & Anderson, T. R. MEDUSA-2.0: an intermediate complexity biogeochemical model of the marine carbon cycle for climate change and ocean acidification studies. Geosci. Model Dev. 6, 1767–1811 (2013).

19. Tittensor, D. P. et al. A protocol for the intercomparison of marine fishery and ecosystem models: Fish-MIP v1. 0. Geosci. Model Dev. 11, 1421–1442 (2018).

20. Lotze, H. K., Tittensor, D. P., Bryndum-buchholz, A., Eddy, T. D. & Cheung, W. W. L. Global ensemble projections reveal trophic amplification of ocean biomass declines with climate change. 116, (2019).

21. Kroodsma, D. A. et al. Tracking the global footprint of fisheries. Science (80-.). 359, 904–908 (2018).

22. Behrenfeld, M. J. & Falkowski, P. G. Photosynthetic rates derived from satellite-based chlorophyll concentration. Limnol. Oceanogr. 42, 1–20 (1997).

23. Haug, T. et al. Future harvest of living resources in the Arctic Ocean north of the Nordic and Barents Seas: A review of possibilities and constraints. Fish. Res. 188, 38–57 (2017).

24. Staresinic, N., Farrington, J., Gagosian, R. B., Clifford, C. H. & Hulburt, E. M. Downward transport of particulate matter in the Peru coastal upwelling: role of the anchoveta, Engraulis ringens. in Coastal Upwelling Its Sediment Record 225–240 (Springer, 1983).

25. Bisson, K., Siegel, D. A. & DeVries, T. Diagnosing mechanisms of ocean carbon export in a satellite-based food web model. Front. Mar. Sci. (2020).

26. Casini, M. et al. Multi-level trophic cascades in a heavily exploited open marine ecosystem. Proc. R. Soc. B Biol. Sci. 1793–1801 (2008). doi:10.1098/rspb.2007.1752

27. Duarte, C. M. et al. Rebuilding marine life. Nature 580, 39–51 (2020).

28. Pusceddu, A. et al. Chronic and intensive bottom trawling impairs deep-sea biodiversity and ecosystem functioning. Proc. Natl. Acad. Sci. 111, 8861 LP – 8866 (2014).

29. Schmidt, K. et al. Zooplankton Gut Passage Mobilizes Lithogenic Iron for Ocean Productivity. Curr. Biol. 26, 2667–2673 (2016).

30. Ratnarajah, L., Bowie, A. R., Lannuzel, D. & Klaus, M. The Biogeochemical Role of Baleen Whales and Krill in Southern Ocean Nutrient Cycling. 1–18 (2014). doi:10.1371/journal.pone.0114067

31. Shatova, O., Wing, S. R., Gault-Ringold, M., Wing, L. & Hoffmann, L. J. Seabird guano enhances phytoplankton production in the Southern Ocean. J. Exp. Mar. Bio. Ecol. 483, 74–87 (2016).

32. Votier, S. C. et al. Changes in fisheries discard rates and seabird communities. Nature 427, 727–730 (2004).

33. Laufkötter, C. et al. Projected decreases in future marine export production: the role of the carbon flux through the upper ocean ecosystem. Biogeosciences 13, 4023–4047 (2016).

34. Laufkötter, C. et al. Drivers and uncertainties of future global marine primary production in marine ecosystem models. Biogeosciences 12, 6955–6984 (2015).

35. Möllmann, C., Müller-Karulis, B., Kornilovs, G. & St John, M. A. Effects of climate and overfishing on zooplankton dynamics and ecosystem structure: regime shifts, trophic cascade, and feedback loops in a simple ecosystem. ICES J. Mar. Sci. 65, 302–310 (2008).

36. Poloczanska, E. S. et al. Responses of Marine Organisms to Climate Change across Oceans. Frontiers in Marine Science 3, 62 (2016).

37. Bopp, L. et al. Potential impact of climate change on marine export production. Global Biogeochem. Cycles 15, 81–99 (2001).

38. Worm, B. et al. Rebuilding Global Fisheries. Science (80-.). 325, 578 LP – 585 (2009).

39. Murawski, S. A. Rebuilding depleted fish stocks: the good, the bad, and, mostly, the ugly. ICES J. Mar. Sci. 67, 1830–1840 (2010).

40. Long, R. D., Charles, A. & Stephenson, R. L. Key principles of marine ecosystem-based management. Mar. Policy 57, 53–60 (2015).

41. Oregon State University. Science.oregonstate.edu. (2017). Available at: http://www.science.oregonstate.edu/ocean.productivity/index.php. (Accessed: 5th June 2017)

42. Dunne, J. P., Armstrong, R. a., Gnanadesikan, A. & Sarmiento, J. L. Empirical and mechanistic models for the particle export ratio. Global Biogeochem. Cycles 19, GB4026 (2005).

43. Taconet, M., Kroodsma, D. A. & Fernandes, J. A. Global Atlas of AIS-based fishing activity - Challenges and opportunities. (2019).

44. Watson, R. A. A database of global marine commercial, small-scale, illegal and unreported fisheries catch 1950-2014. Sci. data 4, 1–9 (2017).

