## Supplementary Information for "Commercial fishery disturbance of the global open-ocean carbon sink"

Cavan, E. L[1]. & Hill, S. L.[2]

[1]Imperial College London, Silwood Park Campus, Ascot, Berkshire, SL5 7PY, UK.
[2]British Antarctic Survey, Natural Environment Research Council, High Cross, Madingley Rd, Cambridge, CB3 0ET, UK.

**Contents**

1. Supplementary Data, Carbon export and fishing for each fishery area, Table S1
2. Supplementary Methods, Information on catch categories, Table S2
3. Supplementary Methods, Indicators of carbon sink intensity and fishing intensity, Fig. S1
4. Supplementary Discussion text, Evidence for localised dead zones as a result of discarding
5. References

**Table S1**. Sea surface temperature (SST), POC export and fisheries data for each FAO major area. POC export data (Fig. 1a, GtC $yr^{-1}$) and Fishing intensity (Fig. 1b, as hours $x10^{6}$) for each fishery area summed. We also present fishery effort as a percentage of fisheries catch globally. Main gear types and fished groups cumulatively contributing to ≥ 50 % of catch are reported, their contribution (%) is given in parentheses. For gear type T = trawl, PS = purse seine, D = dredge, SG = set gillnet, LL = Longline, UG = unknown gear. For species GF = ground, SP = small pelagic, LP = large pelagic, DF = deep water and UF = unspecified fish, PC = pelagic, BC = benthic and UC = unspecified crustaceans, UM = unspecified molluscs, B = bivalves and S = squid. *Gear type from FAO global capture production data rather than landings data.

| FAO Area | Name | Mean SST (ºC) | Total POC export, GtC $yr^{-1}$ (% globally) | Total fishing intensity, hours x10^6 (% globally) | Fisheries catch, % globally | Main gear type (% catch) | Main fished groups (% catch) |
|---|---|---|---|---|---|---|---|
| 18 | Arctic Sea | 01.7 | 0.24 (7 %) | 0.29 (0 %) | 0 % | T (36), PS (32) | GF (85) |
| 21 | NW Atlantic | 11.3 | 0.15 (4 %) | 8.16 (4 %) | 2 % | D (31), PT (23) | PC (13), B (12), SP (21), BC (7) |
| 27 | NE Atlantic | 08.9 | 0.46 (14 %) | 33.26 (15 %) | 11 % | T (72)* | SP (38), GF (13) |
| 31 | Central W Atlantic | 26.6 | 0.07 (2 %) | 4.4 (2 %) | 2 % | PS (46), T (23) | SP (36), UF (10), B (5) |
| 34 | Central E Atlantic | 25.0 | 0.11 (3 %) | 14.02 (6 %) | 6 % | PS (39), T (30) | SP (54) |
| 37 | Mediterranean | 22.6 | 0.03 (1 %) | 10.22 (5 %) | 2 % | T (38), PS (29) | SP (44), UF (4), B (4) |
| 41 | SW Atlantic | 19.1 | 0.21 (6 %) | 10.16 (5 %) | 2 % | T (70) | S (21), GF (22), BC (5), DF (4) |
| 47 | SE Atlantic | 18.4 | 0.20 (6 %) | 7.27 (3 %) | 2 % | T (39), PS (31) | SP (34), GF (19) |
| 48 | Antarctic Atlantic | 0.40 | 0.09 (3 %) | 0.39 (0 %) | 0 % | T (99) | PC (99) |
| 51 | W Indian | 24.6 | 0.19 (5 %) | 18.9 (9 %) | 6 % | T (38), SG (22) | UF (14), SP (13), LP (12), GF (8), DF (3), PC (3) |
| 57 | E Indian | 19.5 | 0.22 (6 %) | 5.91 (3 %) | 8 % | T (33), SG (31) | UF (30), SP (12), LP (3), PC (2), UC (2) GF (2) |
| 58 | Antarctic Indian | 01.9 | 0.10 (3 %) | 0.53 (0 %) | 0 % | LL (95) | DF (79) |
| 61 | NW Pacific | 18.5 | 0.32 (9 %) | 29.99 (14 %) | 27 % | T (49), UG (13) | GF (15), SP (14), UF (13), UM (3), PC (3), S (2), LP (2) |
| 67 | NE Pacific | 11.6 | 0.20 (6 %) | 5.53 (3 %) | 4 % | T (77) | GF (54) |
| 71 | Central W Pacific | 28.7 | 0.10 (3 %) | 29.4 (13 %) | 15 % | T (45), PS (13) | UF (23), LP (17), SP (8), S (2) |
| 77 | Central E Pacific | 25.8 | 0.18 (5 %) | 26.69 (12 %) | 2 % | PS (56) | SP (42), LP (9) |
| 81 | SW Pacific | 14.3 | 0.23 (7 %) | 6.08 (3 %) | 1 % | PS (58) | DF (24), UF (11), SP (14), S (5) |
| 87 | SE Pacific | 18.5 | 0.24 (7 %) | 9.17 (4 %) | 12 % | PS (78) | SP (57) |
| 88 | Antarctic Pacific | 00.3 | 0.06 (2 %) | 0.12 (0 %) | 0 % | LL (100) | DF (95) |

Information on catch categories

The FAO Global Capture Production database[1] provides catch information by taxon. In most cases taxon is resolved to species, but in some cases it is resolved to a lower taxonomic level (e.g. "sardinellas" or "various squids"). We identified the taxa which dominate the catch in each fishing area (i.e. the top ranking taxa in terms of catch weight, which constitute 50 % or the closest value above 50 % of the overall catch). This resulted in a list of 62 species which we assigned to one of ten categories of catch species (Table S2) on the basis of the following definitions:

Small pelagic fish (SP); pelagic fish with maximum length < 60cm

Large pelagic fish (LF); pelagic fish with maximum length > 60cm

Groundfish (G); all shelf-associated demersal fish

Deep water fish (DF); bottom-associated fish found in off shelf waters (typically deeper than 500 m)

Unspecified fish (UF);

Pelagic crustaceans (PC);

Benthic crustaceans (BC); all bottom-associated crustaceans

Unspecified crustaceans (UC);

Unspecified molluscs (UC);

Squid (S); all squid

Bivalves (B); all bivalves

**Table S2**. Classification of dominant taxa recorded in the FAO Global Capture Production database[1] into the catch categories used in the current study.

| Taxa | Category | Abbreviation |
|---|---|---|
| American lobster | Benthic crustacean | BC |
| Argentine red shrimp | Benthic crustacean | BC |
| Marine crustaceans nei | Unspecified crustaceans | UC |
| Antarctic toothfish | Deep water fish | DF |
| Blue grenadier | Deep water fish | DF |
| Bombay-duck | Deep water fish | DF |
| Patagonian grenadier | Deep water fish | DF |
| Patagonian toothfish | Deep water fish | DF |
| Alaska pollock(=Walleye poll.) | Groundfish | G |
| Argentine hake | Groundfish | G |
| Atlantic cod | Groundfish | G |
| Cape hakes | Groundfish | G |
| Croakers, drums nei | Groundfish | G |
| Hairtails, scabbardfishes nei | Groundfish | G |
| Largehead hairtail | Groundfish | G |
| Pacific cod | Groundfish | G |

| | | |
|---|---|---|
| Whitemouth croaker | Groundfish | G |
| Seerfishes nei | Large pelagic fish | LP |
| Skipjack tuna | Large pelagic fish | LP |
| Yellowfin tuna | Large pelagic fish | LP |
| Akiami paste shrimp | Pelagic crustacean | PC |
| Antarctic krill | Pelagic crustacean | PC |
| Natantian decapods nei | Pelagic crustacean | PC |
| Northern prawn | Pelagic crustacean | PC |
| Anchoveta(=Peruvian anchovy) | Small pelagic fish | SF |
| Atlantic chub mackerel | Small pelagic fish | SP |
| Atlantic herring | Small pelagic fish | SP |
| Atlantic mackerel | Small pelagic fish | SP |
| Atlantic menhaden | Small pelagic fish | SP |
| Bonga shad | Small pelagic fish | SP |
| California pilchard | Small pelagic fish | SP |
| Cape horse mackerel | Small pelagic fish | SP |
| Capelin | Small pelagic fish | SP |
| Clupeoids nei | Small pelagic fish | SP |
| European anchovy | Small pelagic fish | SP |
| European pilchard(=Sardine) | Small pelagic fish | SP |
| European sprat | Small pelagic fish | SP |
| Gulf menhaden | Small pelagic fish | SP |
| Hilsa shad | Small pelagic fish | SP |
| Indian mackerel | Small pelagic fish | SP |
| Indian mackerels nei | Small pelagic fish | SP |
| Indian oil sardine | Small pelagic fish | SP |
| Jack and horse mackerels nei | Small pelagic fish | SP |
| Japanese anchovy | Small pelagic fish | SP |
| Pacific anchoveta | Small pelagic fish | SP |
| Pacific chub mackerel | Small pelagic fish | SP |
| Pacific thread herring | Small pelagic fish | SP |
| Round sardinella | Small pelagic fish | SP |
| Sardinellas nei | Small pelagic fish | SP |
| Scads nei | Small pelagic fish | SP |
| Short mackerel | Small pelagic fish | SP |
| Southern African anchovy | Small pelagic fish | SP |
| Southern blue whiting | Small pelagic fish | SP |
| Marine fishes nei | Unspecified fish | UF |
| American cupped oyster | Bivalve | B |
| American sea scallop | Bivalve | B |
| Striped venus | Bivalve | B |
| Argentine shortfin squid | Squid | S |
| Cephalopods nei | Squid | S |
| Various squids nei | Squid | S |
| Wellington flying squid | Squid | S |
| Marine molluscs nei | Unspecified molluscs | UM |

Indicators of carbon sink intensity and fishing intensity

There are no data available that provide reliable measures of either the ultimate benefit of the open-ocean carbon sink (the rate of deep carbon sequestration) or the ecosystem impact of fishing at the spatial resolution of our analysis. We therefore used available indicator variables. Here we briefly discuss the relationship between our indicator variables and the variables that they indicate.

The phytoplankton-driven carbon sink initially depends on the amount of primary production in the euphotic zone, and the proportion of this which gets exported out of the euphotic zone (i.e. export efficiency). As explained in the Methods there is a consensus about how export efficiency varies across the global oceans, and hence we deem this the most appropriate metric of the potential carbon sink on a global scale. However, exported sinking carbon then attenuates rapidly through the mesopelagic zone, being consumed by heterotrophs such as zooplankton and bacteria, meaning only ~ 1 % reaches the deep ocean (i.e. > permanent thermocline)[2,3]. How this varies across the global oceans is not clear, with some meta-analyses suggesting an inverse relationship with temperature[4] and others a positive relationship[5]. In addition, the varying depth of the permanent thermocline needs to be considered[6]. To realise global maps of deep ocean phytoplankton-derived carbon sequestration more needs to be understood on the controls of mesopelagic carbon attenuation[7].

Some of the identified routes of fishery disturbance to the carbon sink (such as removing low-mid trophic level pellet-producing species) are directly related to catch (i.e. the amount of biomass removed). Others (e.g. sediment disturbance) are more directly related to effort (i.e. the amount of time spent fishing). Global effort data are available at the spatial resolution of our analysis (Fig. 2) whereas available catch data are generally aggregated at coarser spatial scales, including the FAO major fishing areas. The catch achieved per unit effort depends on the local biomass of fish and the efficiency of the fishing gear. Effort is therefore an imperfect indicator of catch. Nonetheless it is well correlated with catch at the scale of major fishing areas (Fig S1). An important caveat with the effort data is that the ecosystem impact per unit fishing time varies with multiple factors including gear type (which is considered in our discussion), size of vessel, and other technologies used (e.g. fish finders and fish aggregation devices). Thus our analysis, based on effort data, provides an indication, rather than a definitive identification, of the potential spatial distribution of fishery impacts on the carbon sink and areas of high risk.

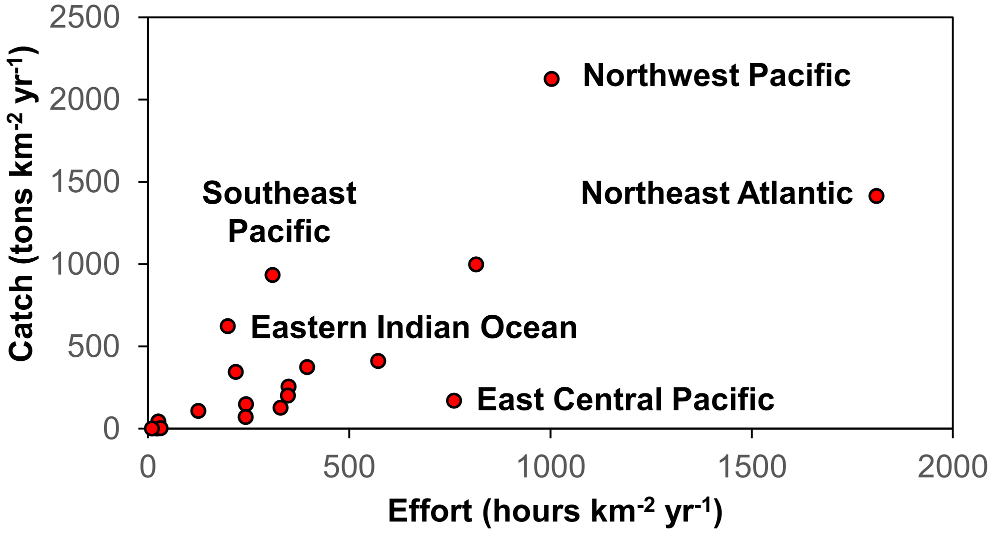


**Fig S1** The relationship between our indicator of fishing impact (effort) and direct impact on target species biomass (catch) at the scale of FAO major fishing areas (Pearson correlation coefficient=0.75, P<0.001), with key outliers labelled. The non-area-adjusted data (catch in tons $yr^{-1}$ versus effort in hours $yr^{-1}$) are also correlated (Pearson correlation coefficient=0.72, P<0.001).

Evidence for localised dead zones as a result of discarding

We are not aware of any direct evidence that the release of discards (unwanted catch and offal) from fishing vessels causes dead zones (i.e. areas with extremely low levels of dissolved oxygen that can cause mass mortality of most metazoan groups)[8]. Nonetheless oxygen depletion resulting from the decomposition of fisheries discards has previously been identified as a potential impact of discarding[9]. The major cause of marine dead zones is an enhanced flow of organic material to the seabed which increases microbial respiration[8]. Localised increases in benthic microbial respiration have been observed as a result of faeces and uneaten food from salmon cages accumulating on the seabed[10]. Similarly, the accumulation of gelatinous carcasses as a result of mass die-offs has led to localised changes in oxygen demands and a switch from autotrophic to heterotrophic systems[11]. It is plausible that the accumulation of such matter on the seabed as a result of discarding would have a similar effect, although this effect is likely to be both ephemeral and spatially limited.

References


1. FAO. *FAO yearbook. Fishery and Aquaculture Statistics 2017.* (2019).
2. Antia, A. N. *et al.* Basin-wide particulate carbon flux in the Atlantic Ocean: Regional export patterns and potential for atmospheric CO2 sequestration. *Global Biogeochem. Cycles* **15**, 845–862 (2001).
3. Turner, J. T. Zooplankton fecal pellets, marine snow, phytodetritus and the ocean's biological pump. *Prog. Oceanogr.* **130**, 205–248 (2015).
4. Henson, S. A., Sanders, R. & Madsen, E. Global patterns in efficiency of particulate organic carbon export and transfer to the deep ocean. *Global Biogeochem. Cycles* **26**, n/a-n/a (2012).
5. Marsay, C. *et al.* Attenuation of sinking particulate organic carbon flux through the mesopelagic ocean. *Proc. Natl. Acad. Sci.* **12**, 1089–1094 (2015).
6. Palevsky, H. I. & Doney, S. C. How Choice of Depth Horizon Influences the Estimated Spatial Patterns and Global Magnitude of Ocean Carbon Export Flux. *Geophys. Res. Lett.* **45**, 4171–4179 (2018).
7. Cavan, E. L., Laurenceau-Cornec, E. C., Bressac, M. & Boyd, P. W. Exploring the ecology of the mesopelagic biological pump. *Prog. Oceanogr.* 102125 (2019). doi:https://doi.org/10.1016/j.pocean.2019.102125
8. Diaz, R. J. & Rosenberg, R. Spreading Dead Zones and Consequences for Marine Ecosystems. *Science (80-. ).* **321**, 926 LP – 929 (2008).
9. Clucas, I. *A study of the options for utilization of bycatch and discards from marine capture fisheries. FAO.* (1997).
10. Findlay, R. H., Watling, L. & Mayer, L. M. Environmental impact of salmon net-pen culture on marine benthic communities in Maine: A case study. *Estuaries* **18**, 145 (1995).
11. Guy-Haim, T. *et al.* The effects of decomposing invasive jellyfish on biogeochemical fluxes and microbial dynamics in an ultraoligotrophic sea. *Biogeosciences Discuss.* **2020**, 1–37 (2020).
